# A versatile set of computational tools for inference of genetic ancestry from human molecular data

**DOI:** 10.64898/2026.10.09.758080

**Authors:** Pascal Belleau, Xintong Li, Astrid Deschênes, Laine Marrah, Alexander Krasnitz

## Abstract

Molecular properties of human diseases and conditions often exhibit dependence on the genetic ancestry of the donor. In order to make more human molecular data available for ancestry-informed research, we developed a set of software tools termed Robust Ancestry Inference using Data Synthesis (RAIDS) for inference of genetic ancestry from sequence data that pose challenges to genotyping. The data-synthesis procedure, underlying RAIDS, makes it a unique tool capable of assessing and optimizing the accuracy of inference for each individual molecular profile, with no resort to cohort data. Here we describe a recent expansion and refinement of RAIDS, applicable to nucleic-acid sequence data resultant from a broad range of experimental strategies. These strategies currently include whole-genome, whole-exome, and targeted gene panel sequencing, RNA-Seq, including single-cell RNA-Seq, chromatin-accessibility (ATAC-Seq), and protein-binding (ChIP-Seq) assays. RAIDS inferential tools perform robustly in the presence of somatic nucleic-acid alterations caused by cancer and under the conditions of extreme non-uniformity of sequence read coverage, yielding both global ancestry and ancestral admixtures of the donor. RAIDS is available as an eponymous Bioconductor package.

## Introduction

Genetic ancestry significantly influences human health and disease, affecting phenotypes across diverse conditions including cancer, Alzheimer’s disease, and viral immune responses^1–7^. These genetic effects align with epidemiological evidence linking race and ethnicity to disease incidence, clinical progression, and outcomes^8–15^. In cancer, large-scale genomic characterization reveals ancestry-associated differences in tumor molecular landscapes, including somatic driver mutation frequencies, genomic instability, copy number variation, and immune microenvironments^16–21^. These molecular variations have been observed to fundamentally impact treatment response and the risk of adverse side effects^22,23^.

Discovery of these ancestry-specific mechanisms is severely hampered by a shortage of molecular data from non-European populations. In The Cancer Genome Atlas (TCGA), for instance, certain indigenous populations are critically underrepresented, featuring only 27 self-identified Alaskan Natives/American Indians and 13 Native Hawaiians/Pacific Islanders. Even within larger aggregate cohorts, sample sizes drop precipitously when stratified by cancer type; for example, out of 934 African American patients in TCGA, only 7 cases are available for prostate and pancreatic cancers each.

Historically, ancestry annotation has relied on self-identified race and ethnicity (SIRE). While SIRE captures essential socio-cultural determinants of health, it is frequently omitted, incomplete, or inconsistent with genetic background. A PubMed survey revealed that only 37% of cancer sequencing studies reported race and 17% reported ethnicity^24^, while SIRE is missing from 56% of electronic health records^25^. Furthermore, SIRE relies on broad categories that fail to capture granular genetic backgrounds or quantify ancestral admixtures.

Direct germline DNA genotyping offers a highly accurate alternative. Standard pipelines leverage whole-genome sequencing (WGS), whole-exome sequencing (WES), or microarray data to infer ancestry via population-specific single-nucleotide variants (SNVs)^26–28^, as demonstrated in ~10,000 TCGA donors^18,21^. However, matching germline profiling is frequently omitted in tissue-specific molecular studies or non-cancer phenotypic assays. In these scenarios, genetic ancestry must be inferred from alternative molecular modalities, such as RNA sequencing (RNA-Seq), targeted gene panels, chromatin immunoprecipitation sequencing (ChIP-Seq) or transposase-accessible chromatin using sequencing (ATAC-Seq).

Inferring ancestry from these non-standard datasets introduces unique technical challenges. First, data-specific properties create non-uniformities: RNA-Seq yields highly uneven transcript coverage driven by variable expression levels, protocol design, and allele-specific expression; similar extreme coverage non-uniformities affect ATAC-Seq and ChIP-Seq data. Narrowly targeted panels cover only a very small fraction of the genome. Second, cancer-derived sequences are heavily distorted by somatic alterations—including copy number variants, translocations, loss of heterozygosity, microsatellite instabilities, and somatic SNVs—all of which obstruct accurate ancestry inference. Consequently, the accuracy of ancestry inference from such data is uncertain and may vary widely from one profile to another. The inference accuracy may be gauged in a large cohort, where accurate ancestry calls from matching germline DNA sequences, e.g., deep whole-genome sequences, are available for comparison. Often, neither is available, and even when it is, accuracy estimated for a cohort need not apply to an individual profile with distinct sequence properties. We therefore face two critical questions:

- **A:** How can the accuracy of genetic ancestry inference be assessed and optimized for a given molecular profile’s unique sequence properties (e.g., coverage uniformity, depth, read length, and quality)?
- **B:** How can this be achieved without sequencing data for a large, ancestrally diverse control cohort, with the underlying biology and technical properties similar to those of the target profile?

Little systematic work has addressed ancestry inference from inputs other than germline DNA. Prior to our work^29^ studies have been limited to RNA-Seq, deeply sequenced gene panels and cancer-derived data from SNP microarrays, whole exomes and whole genomes^17,30–33^. For all such studies, benchmarking, if at all, was performed against ancestry inferred from matching germline DNA. To our knowledge, no prior work has inferred ancestry from chromatin-profiling sequences, in particular, ATAC or ChIP sequences.

Addressing the two critical questions as posed requires a systematic computational framework capable of ancestry inference, alongside an adaptive mechanism to optimize and assess accuracy based on data quality and coverage depth of any given individual sequence profile. We recently developed such a framework. It combines existing inference algorithms with a procedure we termed “data synthesis”, which answers both critical questions: it assesses performance and optimizes inference parameters for each individual profile, using synthetic profiles that retain the sequence properties of that profile, so that no ancestrally diverse control cohort of similar data is required. Our methodology, termed RAIDS (Robust Ancestry Inference using Data Synthesis), has demonstrated consistently high accuracy across ancestral groups, cancer types and sequencing modalities, including WES, deeply sequenced gene panels and RNA-Seq^29^. RAIDS has been implemented and released as an eponymous R/Bioconductor package^34,35^.

Here, we report a major functional expansion of *RAIDS*. First, we extend its inferential capability to sequences of single-cell RNA (scRNA-Seq) and those produced using chromatin-probing experimental strategies, namely, ATAC-Seq and ChIP-Seq. Second, we extend the capabilities of *RAIDS* to accurate and robust inference of ancestral admixtures across multiple key molecular protocols. Given the methods-development focus of this work, we present some of the key points of the methodology underlying RAIDS in the Results section and fill in further methodological details in the Methods section.

## Results

### RAIDS is applicable to molecular data from experimental strategies targeting chromatin and single-cell RNA

In the preceding publication we demonstrated applicability of RAIDS to human molecular data from whole-exome, RNA-Seq and narrowly-targeted gene panel sequences. For all these experimental strategies, the applicability extended to sequences of nucleic acids affected by cancer. We validated these findings for 16 representative cancer types. Here we report the applicability of RAIDS to sequences from additional experimental protocols, namely, single-cell RNA sequencing and two widely used chromatin profiling methods, ATAC-Seq and ChIP-Seq for 4 ChIP-Seq targets. We note that scRNA sequencing, as a rule, does not yield sequences with coverage sufficient for inference of genetic ancestry from RNA sequences of individual cells. Instead, genetic ancestry is inferred from an aggregate sequence of multiple cells. The resulting aggregate sequence provides coverage predominantly for the 3’ or 5’ ends of transcripts rather than full transcripts and thereby differs substantially from that of bulk RNA with an equal number of reads. We therefore here consider single-cell RNA sequencing a distinct molecular strategy.

For these molecular strategies, our initial focus in this work was on inference of global continental ancestry as a categorical variable whose possible values are African (AFR), American (AMR), East Asian (EAS), European (EUR) and South Asian (SAS). The input molecular data for all analyses presented here are summarized in Figure 1A, for multiple combinations of molecular strategies. Summaries of the input data for each input dataset are provided in Table S1 and a full, case-by-case tabulation of all the input molecular data is in Table S2. We also inferred, for each human subject, genetic ancestry as a category from the matching WES or RNA-Seq profile. Both of these molecular strategies we previously demonstrated to result in input data for which RAIDS yields accurately inferred ancestry^29^, even in presence of major somatic alterations, such as those caused by cancer. Ancestral composition of all the molecular data used in the present work is summarized in Figure 1B, and in more detail in Tables S2 and S3.

**Figure 1:**
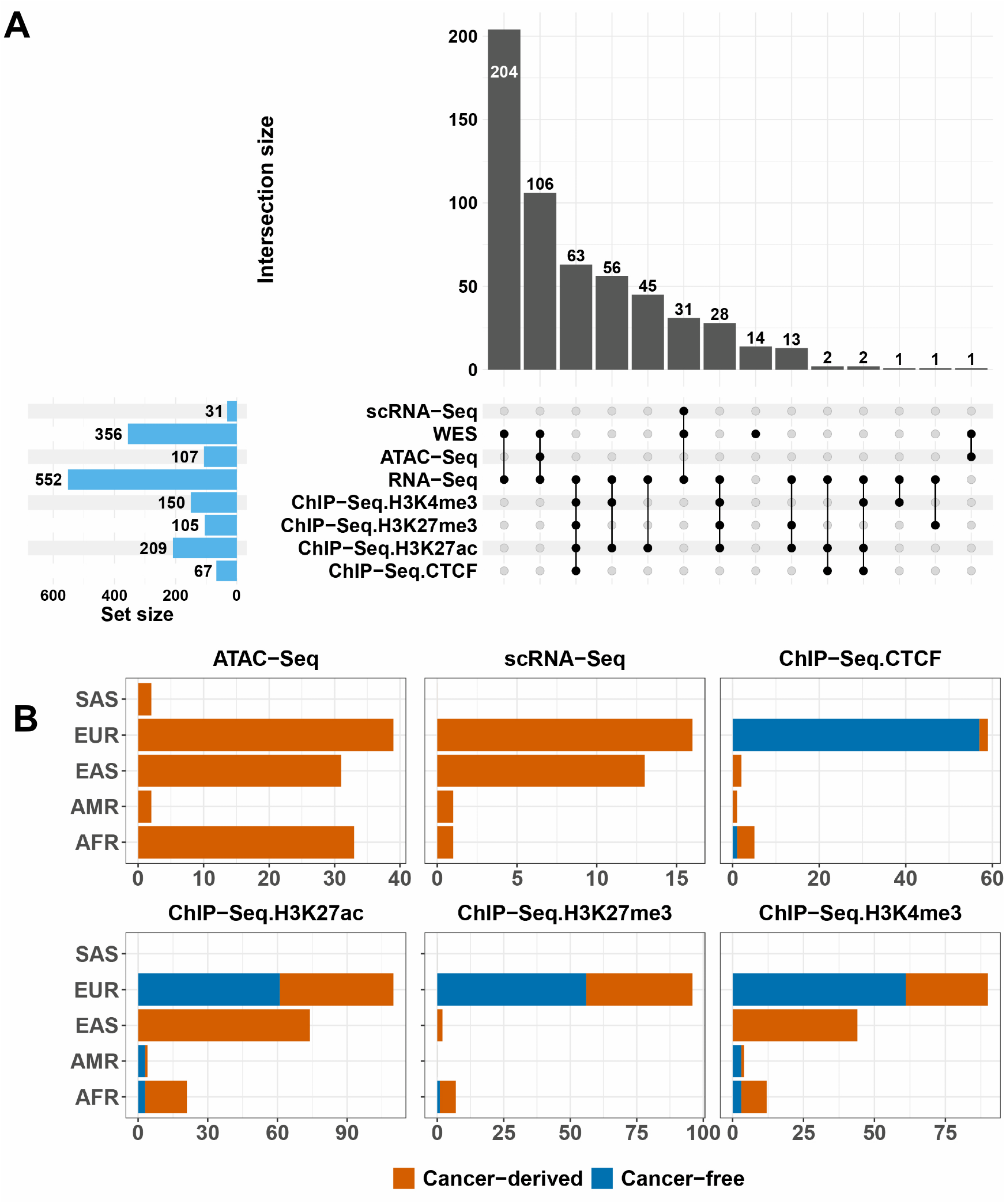
A – summary of the 567 human donor molecular data included in the study, categorized by molecular strategy combinations; B – for each molecular strategy studied, continental ancestral composition of the combined cohort, as determined using matching WES data, where available and RNA-Seq data otherwise, with cancer or cancer-free origin of the data indicated by colors.

In assessing the performance of inference using data synthesis, we followed a procedure established in our prior work, and computed, for each molecular profile and each molecular strategy under study, two measures of performance. These are the accuracy, defined as the observed frequency of correct inference, and the area under the receiver operating characteristic curve (AUROC), both overall and ancestry-specific, for each of the five continental ancestries. As discussed in our prior work^29,35^, these measures depend on the inference parameters. As with the experimental strategies we studied previously, we employed here principal component analysis in combination with K nearest-neighbor classification^27^. Thereby, the inference parameters in this case are the number D of principal dimensions and the number K of nearest neighbors. We determine, for each input profile, a range of parameters yielding optimal or near-optimal performance. This aspect of the inference performance, assessed using data synthesis, is illustrated in Figure 2A, with the underlying data in Table S4, for a small number of randomly chosen nucleic acid donors (for all donors, Tables S5 and S6), and summarized in Figure 2B and Table S7, for all donors with data for a given molecular strategy.

**Figure 2:**
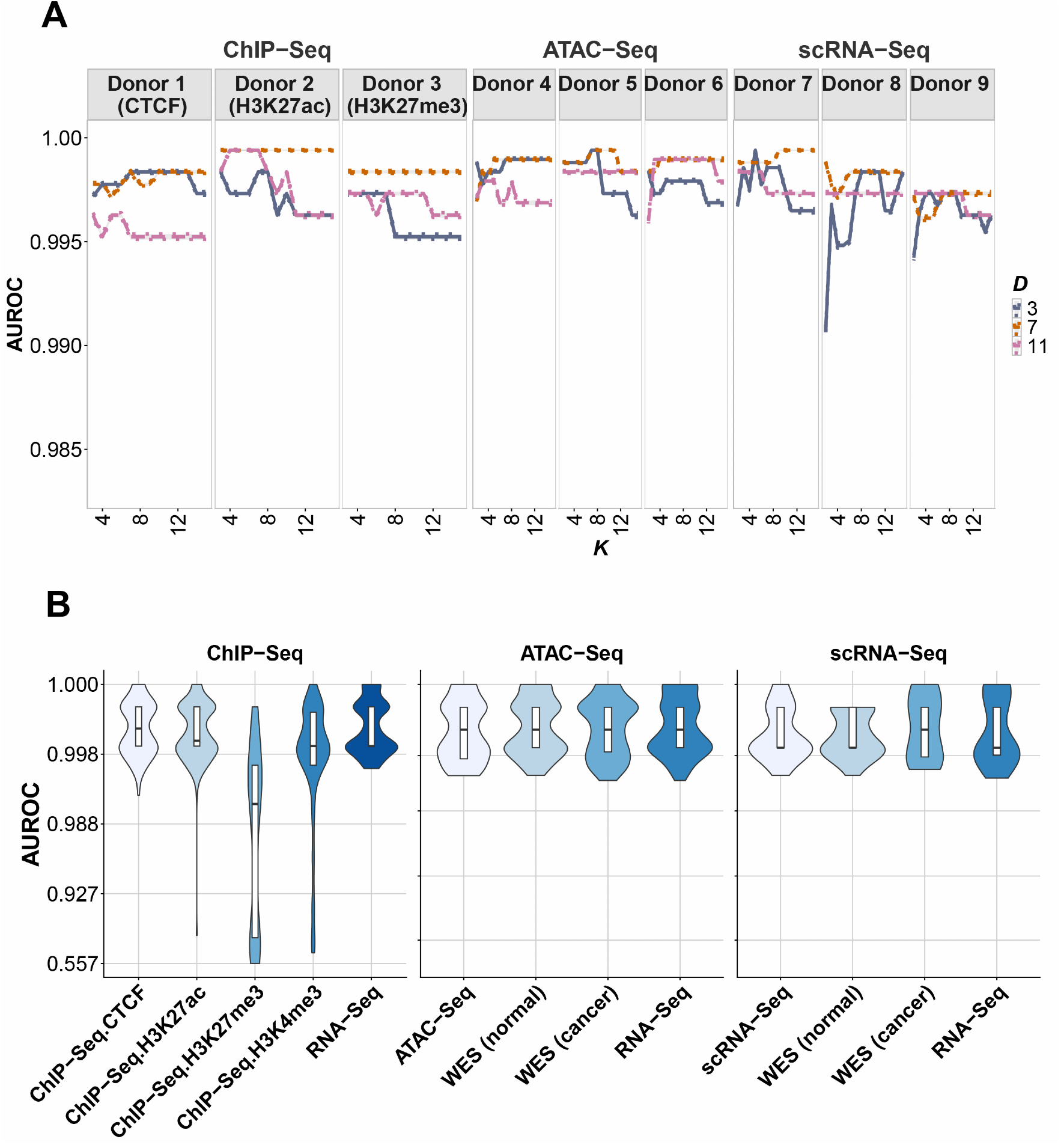
A – performance of RAIDS in inference of ancestry as a category at a continental level of resolution, quantified as area under the ROC curve (AUROC) using data synthesis, as a function of the inference parameters D and K and illustrated, for individual nucleic-acid donors for data from ChIP-Seq (split by ChIP targets), ATAC-Seq and scRNA-Seq; B – cohort-wide summaries, of AUROC, estimated, for each nucleic-acid donor, using data synthesis, across experimental strategies as indicated.

We next quantified cohort-wide performance of RAIDS for scRNA-Seq, ATAC-Seq and ChIP-Seq derived molecular data. To this end, we considered our findings from matching WES or RNA-Seq (for inference from ChIP-Seq profiles) data as gold standard. Cohort-wide performance of RAIDS for these experimental strategies is summarized in Table 1, alongside with the confidence intervals for each measure of performance. As these tabular summaries make clear, RAIDS achieves high accuracy across all the strategies studied. We further note broad consistency between performance assessments from data synthesis and those cohort-wide.

**Table 1:** Cohort-wide measures of performance for inference of ancestry as a category for the molecular data studied.

| Experimental strategy | AUROC | 95% CI | Accuracy | 95% CI2 |
|---|---|---|---|---|
| WES | 0.9957 | 0.9955-0.9959 | 0.9854 | 0.9483-0.9982 |
| RNA-Seq | 0.9500 | 0.9482-0.9518 | 0.9927 | 0.9600-0.9998 |
| ATAC-Seq | 1.0000 | 1.0000-1.0000 | 1.0000 | 0.9661-1.0000 |
| ChIP-CTCF | 0.9167 | 0.9078-0.9255 | 0.9851 | 0.9196-0.9996 |
| ChIP-H3K27ac | 0.9603 | 0.9598-0.9608 | 0.9904 | 0.9659-0.9988 |
| ChIP-H3K27me3 | 0.9107 | 0.909-0.9124 | 0.8000 | 0.7107-0.8717 |
| ChIP-H3K4me3 | 0.9531 | 0.9525-0.9537 | 0.9467 | 0.8976-0.9767 |
| scRNA-Seq | 0.9167 | 0.9059-0.9275 | 0.9677 | 0.8330-0.9992 |

### Inference of ancestral admixtures with RAIDS

The RAIDS computational framework readily accommodates inference of ancestral admixture by running the ADMIXTURE^26^ algorithm in supervised mode, which requires allele frequencies for each genetic ancestral group as input. To obtain these frequencies, we used the 1000 Genomes Project (1KG) reference dataset. By running ADMIXTURE in unsupervised mode on the 1KG donors, we obtained an optimal partition into five major ancestral groups, each characterized by group-specific variant frequencies. These ancestral groups overlap closely, but not entirely, with the four out of five continental populations defined by the 1KG consortium (Figure S1). As expected, the American continental population presents a strong admixture of agEUR and agAMR groups. We also estimated the ancestral admixture proportions of each 1KG donor. For each ancestral group, we then retained a subset of approximately non-admixed (ANA) donors, defined here as those with an admixture proportion of at least 95% for that group. RAIDS framework used these ANA donors to estimate allele frequencies for each genetic ancestral group, and supplied the allele frequencies matrix to ADMIXTURE in supervised mode. The collection of retained non-admixed donors is indicated in the RAIDS population reference data^36^. Further details of this procedure are presented in the Methods section.

To infer the admixtures of an unrelated molecular profile, we follow our previously described procedure^29,35^ to derive a profile-specific subset of positions with high-frequency variants in linkage equilibrium in the 1KG collection, where the genotypes can be confidently established for the profile. We then employ the ADMIXTURE algorithm in supervised mode, with the allele frequency matrix estimated from non-admixed 1KG profiles described earlier.

The performance of the admixture inference in RAIDS is presented here for the following molecular strategies: WGS, WES, RNA-Seq including scRNA-Seq, and ATAC-Seq. For individual molecular profiles, we assessed performance using data synthesis, as follows. The admixture proportions are analogous to probabilities: both are non-negative and add up to 1. This analogy suggests using divergence measures for probability distributions for assessment of performance. We therefore employed the Jensen-Shannon divergence (JSD)^37–39^ to quantify the deviation between the proportions inferred, respectively, from the synthetic molecular profile and the underlying known reference genotypes^40^. We applied this assessment to 1190 molecular profiles of 356 individual nucleic-acid donors from public data collections, as indicated in Table S3. Each nucleic-acid donor has a WES cancer-free profile and at least one tumor-derived sample for a specific experimental strategy (WES, RNA-Seq, scRNA-Seq, and ATAC-Seq). For each, we produced 780 synthetic profiles and computed the corresponding 780 admixture proportions and JSD values. This procedure is explained in more detail in the Materials and Methods section.

First, we restricted our analysis to a subset of 106 nucleic-acid donors (subset A), out of the total of 356 analyzed, for which molecular data for all 3 experimental strategies (WES, RNA-Seq, ATAC-Seq) for the tumor-derived and WES cancer-free profile were available in the admixture analysis. These cases are indicated in Table S3 in the “subset A” column, with a summary by cancer type in Table S8. The resulting admixture proportions for all 780 synthetic profiles are shown in Figure 3A, for a single breast cancer case across the molecular strategies considered here. In Figure 3B, the families of JSD cumulative for all 780 synthetic profiles are shown in Figure 3A, for a single breast cancer case across the molecular strategies considered here. In Figure 3B, the families of JSD cumulative distribution curves, one curve for each cancer case, are depicted. These distributions are to be compared to the distributions of JSD for pairs of synthetic profiles and genotypes of unrelated 1KG individuals, also shown in Figure 3B. As expected, the former family of distributions favors much smaller JSD values than the latter one. Summarizing our findings for the subset A considered, violin plots for mean JSD values estimated by data synthesis for each cancer case are shown in Figure 3C for all molecular strategies considered here.

**Figure 3:**
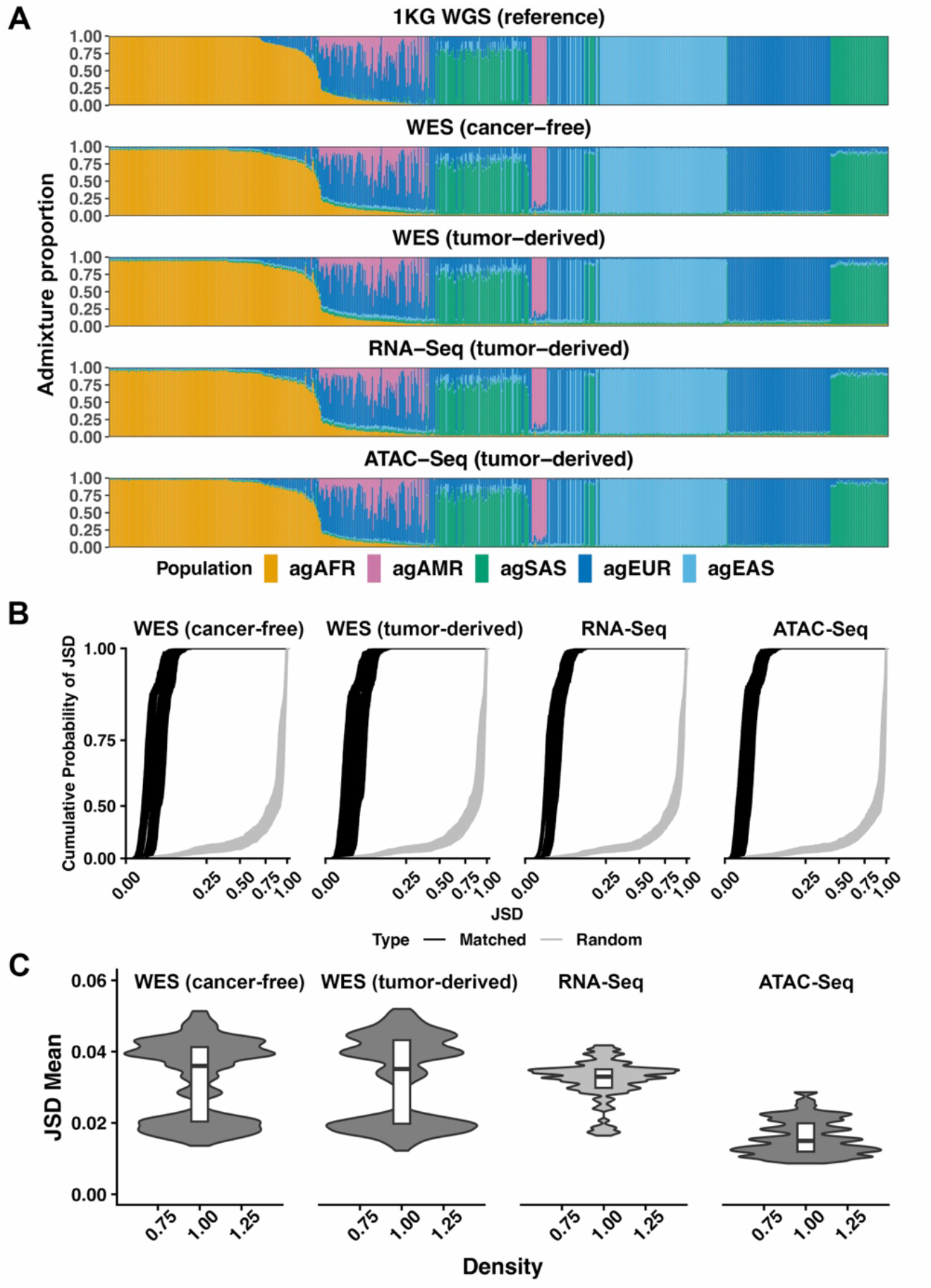
A – ancestral proportions inferred from 780 synthetic profiles for one case of breast cancer, for the patient’s molecular profiles, resulting from experimental strategies as indicated. B – a family of cumulative distributions of the mean JSD measure of performance, as estimated by data synthesis for each of the 106 nucleic-acid donors (subset A), across the four combinations of molecular strategies and nucleic-acid sources as indicated. The corresponding families of distributions of JSD computed for random pairs of genotypes are shown in grey for comparison. C – violin plots for JSD means corresponding each to a distribution shown in B.

Next, we examine cohort-wide performance of RAIDS as admixture inference tool. Figure 4A shows admixture proportions inferred for the subset A, compared across molecular strategies and tissue sources of nucleic acids, as indicated. This comparison indicates close similarity of all admixture proportions computed for the same nucleic-acid donor. More quantitatively, for all molecular protocols studied, the distributions of JSD with the matching (cancer-free) WES-inferred admixtures clearly favor low values, as shown in Figure 4B. We analyzed, along the same lines, all datasets included in this part of the study (in particular scRNA-Seq profiles) (Figures S2-S5 and Tables S9 and S10).

**Figure 4:**
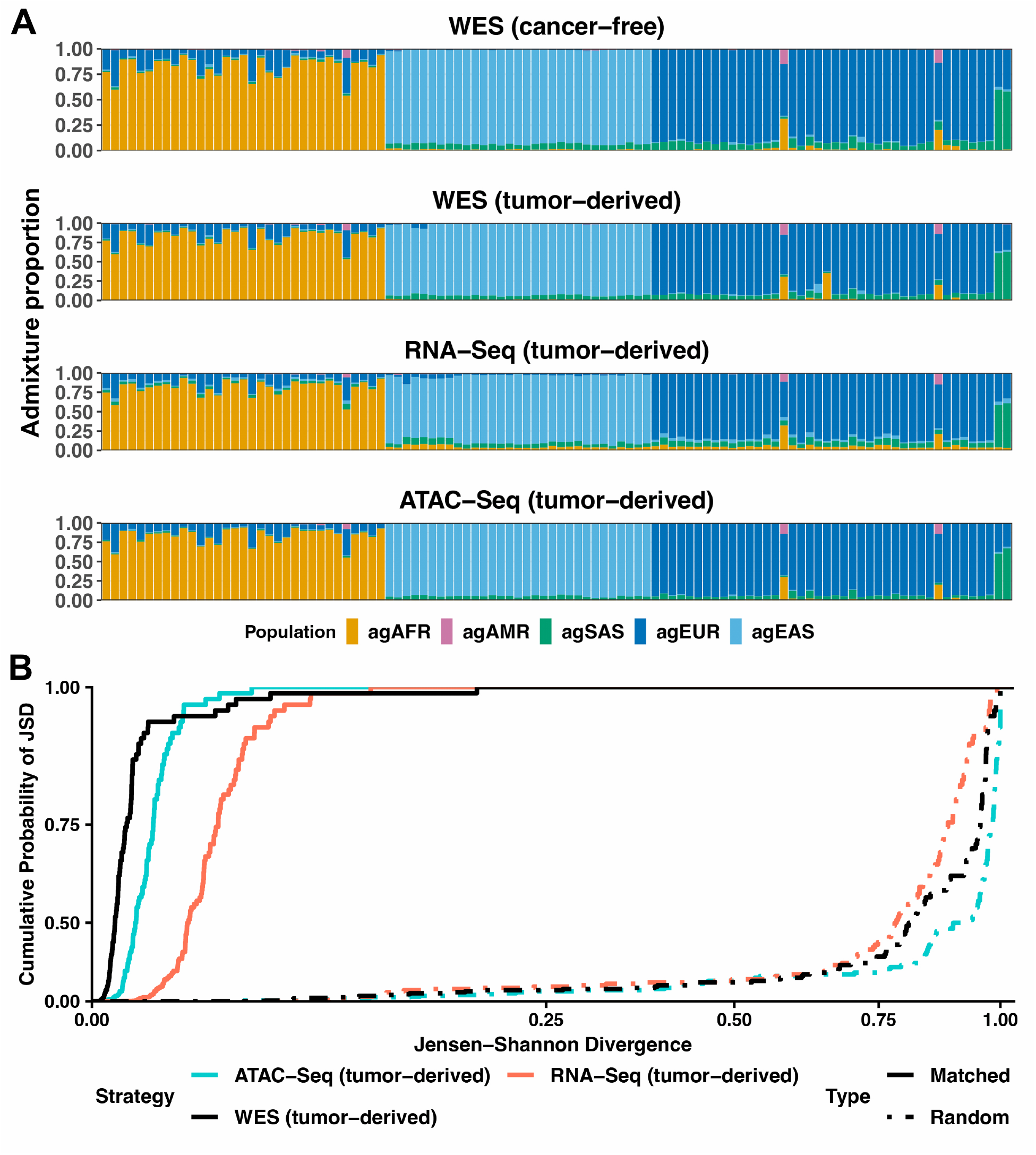
A – cohort-wide admixture proportions for 106 nucleic-acid donors (subset A), across molecular strategies as indicated. B - cumulative distributions of JSD for admixture proportions from cancer-derived molecular data against those derived from matching cancer-free exomes, across molecular strategies as indicated.

## Discussion

The expanded functionalities of RAIDS, as reported here, turn this method into a “Swiss army knife” for inference of genetic ancestry from human molecular data. Data from the molecular strategies that RAIDS can now accommodate account for well over 99% of the human sequence data listed in the Sequence Read Archive (SRA). Thereby, large-scale ancestry-informed analysis of archived or newly produced human sequence data is now possible. Moreover, RAIDS has now been enabled to infer ancestral admixtures across this data spectrum.

The key to the versatility of RAIDS is its core methodology relying on data synthesis, which assesses and optimizes the inference for each individual molecular profile, without a control cohort of similar data. To our knowledge, RAIDS remains the only ancestry-inference tool with this capability. A related strategy was subsequently applied to scRNA-Seq data^41^: inference was evaluated on reference genotypes restricted to the positions genotyped in most donors of a given dataset. This evaluation, however, is performed for a dataset as a whole rather than for an individual profile, requires a cohort of donors with similar data, and does not optimize inference parameters.

Data synthesis, in principle, is applicable to molecular data originating in any sequencing protocol. For this reason, we expect RAIDS to be applicable to low-coverage whole-genome, cytosine-converted and long-read sequences, among others. These expectations will be thoroughly tested and validated by future work. A crucial milestone yet to be reached by RAIDS is the ability to infer local ancestry. This is the present focus of RAIDS development.

The integration of RAIDS into the Bioconductor platform makes it readily available across the research community and facilitates cloud applications of the package. RAIDS project will continue to evolve in lockstep with the nucleic-acid sequencing methodologies.

## Materials and methods

### Population reference dataset

RAIDS requires a population reference dataset for inference of genetic ancestry. As in our earlier publication^29^, the present work relied on the 1000 Genomes (1KG) dataset, integrated into International Genome Sample Resource (IGSR)^42^ and containing genotyping data for 2,469 mutually unrelated individuals. Ancestral composition of the 1KG cohort is detailed in Table S11. The 1KG phased genotyping table was downloaded in the variant call format (VCF)^43^. Continental population designations of all donors — African (AFR), American (AMR), East Asian (EAS), European (EUR), and South Asian (SAS) were also provided by the 1KG Consortium (Table S11).

### Datasets of input molecular profiles for ancestry inference

We collected ChIP-Seq data from the ENCODE and SRA repositories, spanning a broad range of tissues and conditions (Table S2). We confined the analysis to four broadly representative ChIP-Seq targets: the CTCF DNA binding factor, the H3K27ac acetylation mark, and the H3K27me3 and H3K4me3 methylation marks. Matched RNA sequences from all donors in whom ChIP-Seq data originated were used as input into RAIDS to infer the donors’ genetic ancestry, to serve as ground truth for assessment of the RAIDS performance on ChIP sequences. The superpopulation distribution, determined thereby for each ChIP target (Figure 1B) shows representation of all superpopulations, except the South Asian one. All the required ENCODE and SRA sequence data were downloaded in the FASTQ format.

ATAC-Seq, scRNA-Seq input data, alongside the matching RNA and whole-exome sequences, were drawn from several cancer-derived data collections: TCGA, Beat AML1.0^44^, and CPTAC-3^45^. These were obtained from the Genomic Data Commons (GDC). The patients’ SIRE metadata, also provided by the GDC were used to identify subsets of these datasets that are broadly representative of all population groups (Table S3). Ancestral composition of the resultant joint cohort is detailed in Supplementary Table S3 and Figure 1. Sequence data from GDC were downloaded in the BAM format. We note that all cancer-free molecular data from GDC in this study derive from WES.

### Preprocessing of the input molecular profiles and of the population reference dataset

All input molecular profiles in the FASTQ format were mapped to the GRCh38 human reference genome (GDC reference build GRCh38.d1.vd1) using BWA^46^, followed by deduplication and base quality score recalibration with the GATK^47^ utilities. For RNA-Seq data, GATK SplitNCigarReads was executed prior to recalibration. We used these directly for the whole-exome and ATAC sequences, while BAM files containing bulk and single-cell RNA sequences were processed with SplitNCigarReads followed by recalibration.

For all datasets, allele-specific read counts were determined using the snp-pileup utility in the FACETS^48^ package. Reads with a mapping quality score 15 and above and bases with a base quality score above 20 were retained for further analysis. The 1KG population reference data were pre-processed as previously described^29^.

For each input molecular profile to be used as input for inferring genetic ancestry, RAIDS performed, for high-frequency 1KG variant positions, genotype calling, followed by pruning of successfully genotyped positions, to obtain a set of genotypes at positions in linkage equilibrium. For all the downstream analysis, we retained a submatrix of the 1KG genotyping matrix, restricted to positions with high-frequency (> 0.01 in at least one continental population) sequence variants in linkage equilibrium^29,35^. For clarity, Table S12 provides a comprehensive list of the preprocessing software used in this paper, along with respective versions and RRID values.

### Data synthesis

Data synthesis was performed as previously described^29,35^. Specifically, 780 synthetic profiles were produced for each molecular profile of a donor whose ancestry we wish to determine, with underlying known genotypes drawn from the 1KG population reference dataset, 30 for each of the 26 subcontinental populations represented there. The draw from each subcontinental population was at random.

### Inference of genetic ancestry as a category and inference of ancestral admixtures

Inference of ancestry as a category using high-frequency variants in the 1KG genotyping matrix was performed as previously described^29,35^, applying the same algorithm to the additional assay types introduced here (ChIP-Seq, ATAC-Seq, and scRNA-Seq).

For the admixture-proportion analysis, we further processed the 1KG dataset to identify the major ancestral groups represented there and an optimal assignment of the 1KG individuals to these groups. We varied the number G of such groups from 3 through 9. For each G, ADMIXTURE^26^ algorithm with five-fold cross-validation (ADMIXTURE default) was executed five times in the unsupervised mode, each time with a different random-number seed. Figure S6 displays the cross-validation error boxplot throughout this range of G values. From this first step, we retained the interval 3 ≤ G ≤ 5 throughout which the cross-validation error rate deviated little from the optimum. We then chose G = 5, equal to the number of continental populations in the 1KG collection (Figure S1). Using the resulting per-donor admixture proportions, we retained, for each ancestral group, the subset of approximately non-admixed (ANA) donors—defined as those with an admixture proportion of at least 95% in that group across all five random-number generator initializations. RAIDS then estimated the allele frequency matrix (P) using this subset. With this matrix as input, ADMIXTURE was executed in the supervised mode to refine the 1KG admixture proportion matrix^49^. The retained ANA subset of the reference data, as well as the 1KG refined admixture proportion matrix, are included in the preprocessed population reference dataset available with RAIDS^36^.

In order to infer admixtures from a molecular profile, RAIDS executes ADMIXTURE in the supervised mode using the profile genotypes, along with the frequency matrix P from non-admixed profiles in the 1KG collection used to define the ancestral groups.

As in the case of inferring ancestry as a category, data synthesis was used to assess the performance of RAIDS in the inference of ancestral admixtures. Whenever one of the ANA 1KG profiles was used for synthesis, the 1KG genotype frequency matrix P was recomputed with this profile removed, and the admixture proportions of the synthetic profile were inferred using this updated frequency matrix. Strictly, the unsupervised ADMIXTURE analysis used to define the ancestral groups and the ANA subset would also have to be repeated with each such profile removed; we verified that the effect of omitting this step is negligible.

The ancestral admixtures in the synthetic profiles were inferred using ADMIXTURE in the supervised mode, followed by the JSD-based quantification of performance. This procedure is summarized by the following pseudocode, where P[m] denotes the m-th input molecular profile, G[n] the genotypes of the n-th 1KG donor, A[m,n] the admixture proportions inferred from the synthetic profile, T[n] the admixture proportions inferred directly from G[n] and J the JSD of its arguments.

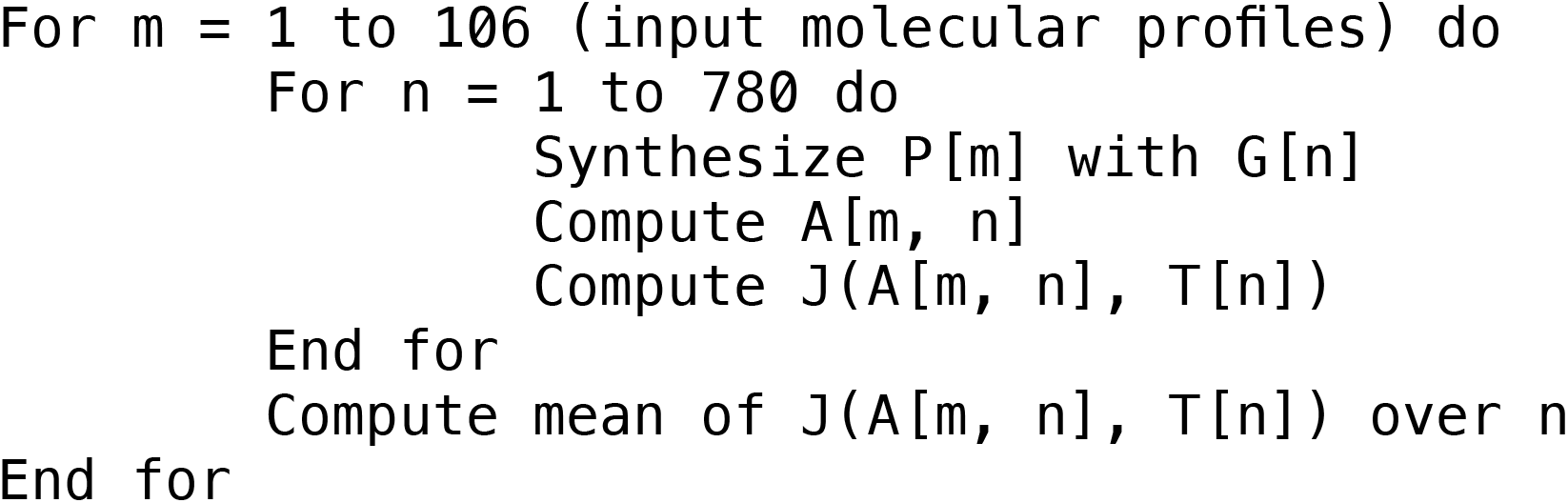

## Acknowledgements

The results presented here are in part based upon data generated by the TCGA Research Network: https://www.cancer.gov/tcga. Also used in this work were data generated by the National Cancer Institute Clinical Proteomic Tumor Analysis Consortium (CPTAC), and the multi-center Beat AML program. This work was supported by a National Institutes of Health award U01 CA289357 to AK.

## Data and code availability

All supplementary tables (S1 through S12) are available at https://labshare.cshl.edu/shares/krasnitzlab/RAIDS2paper. The 1KG reference dataset, preprocessed for use with RAIDS, is available at https://labshare.cshl.edu/shares/krasnitzlab/RAIDSpopRef. The GDC metadata as listed in the supplementary tables are correct as of November 9, 2026. The latest version of RAIDS software package is available as a “devel” option from Bioconductor (https://bioconductor.org/packages/3.24/bioc/html/RAIDS.html, to be included in the official release on November 24, 2026).

