## Supplementary material for "A versatile set of computational tools for inference of genetic ancestry from human molecular data": supplementaryFigures.pdf

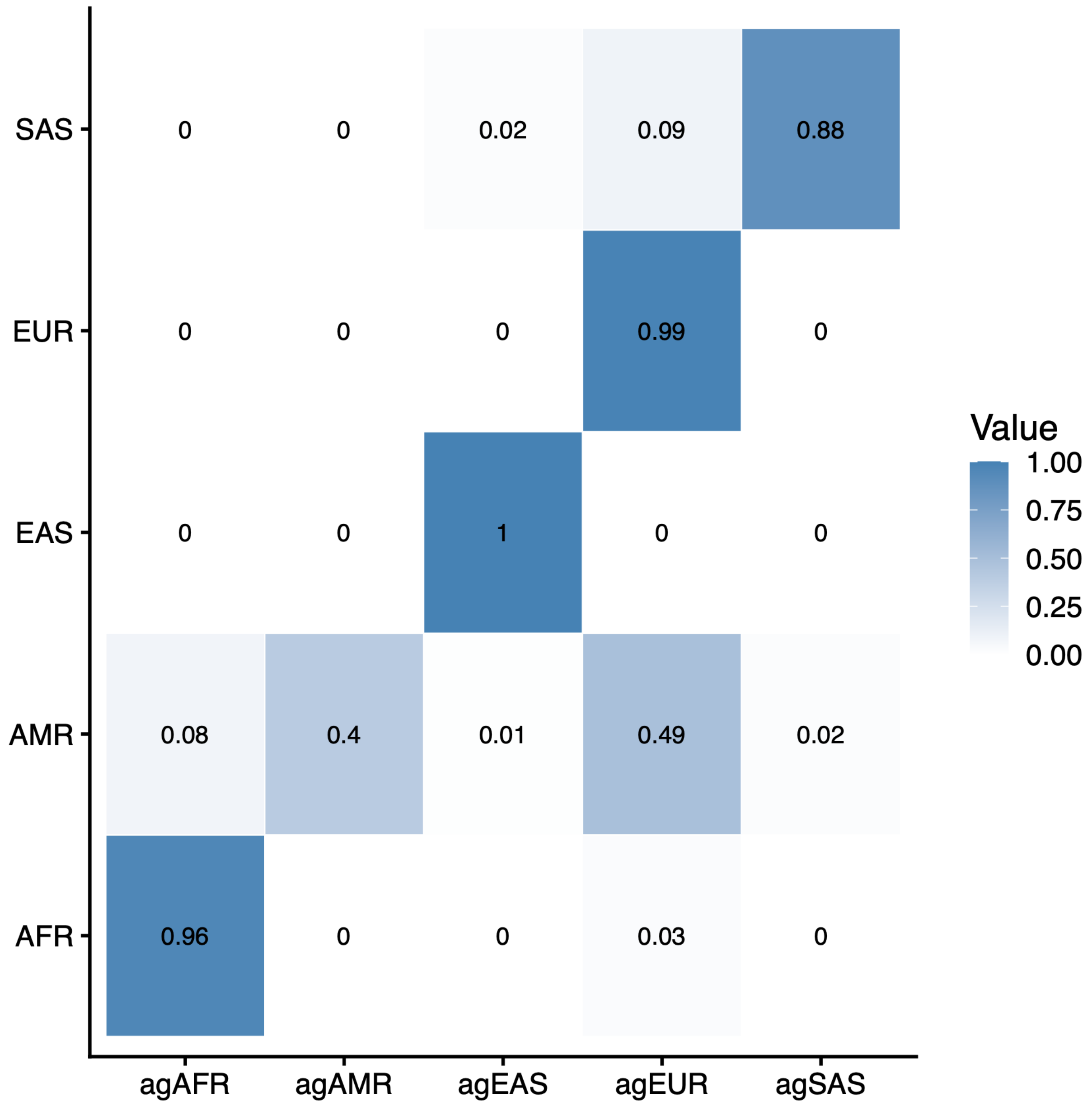


Figure S1: Mean ancestral-group (columns) admixture proportions in 1KG continental-population (rows) samplings.

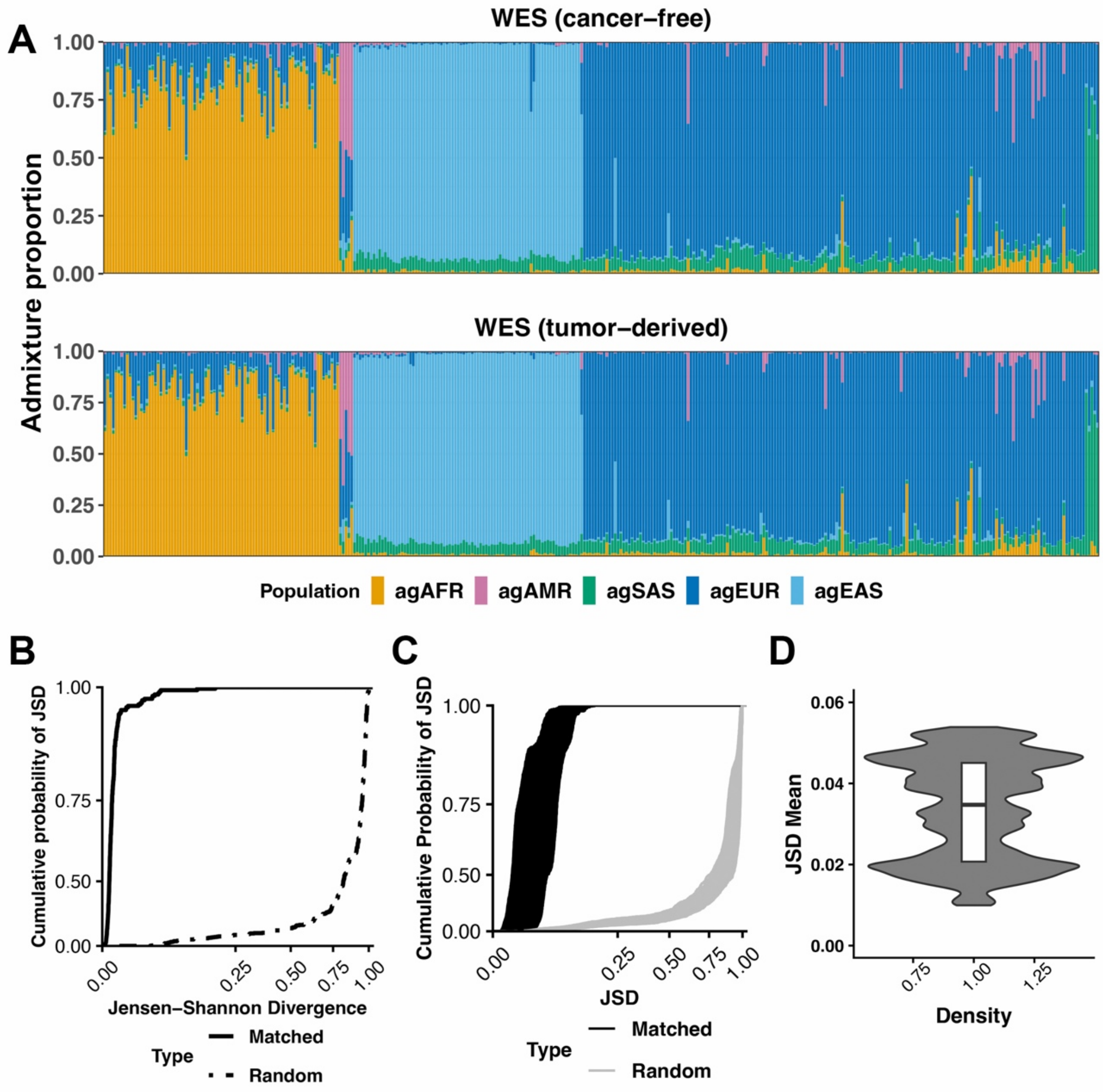


Figure S2: Analysis of admixture proportions, inferred from cancer-derived WES data in a cohort of 355 patient cases representing multiple cancer types, as indicated in Table S3, “Fig. S2 subset” column. A – Cohort-wide comparison of admixture proportions inferred from WES profiles of cancer-free and matching tumor-derived DNA. B – Cohort-wide cumulative distribution of pairwise JSD for admixture proportions inferred from cancer-derived, with those from matching cancer-free whole-exome sequences. Cumulative JSD distribution for random pairs of patients is shown for reference. C – Cohort-wide family of cumulative distributions of the mean JSD measure of performance, estimated, for each patient, by data synthesis for each of the 355 cases of cancer-derived WES. The corresponding family of distributions of JSD computed for random pairs of genotypes is shown for reference. D – Cohort-wide Violin plot for the empirical mean JSD values, each estimated by data synthesis.

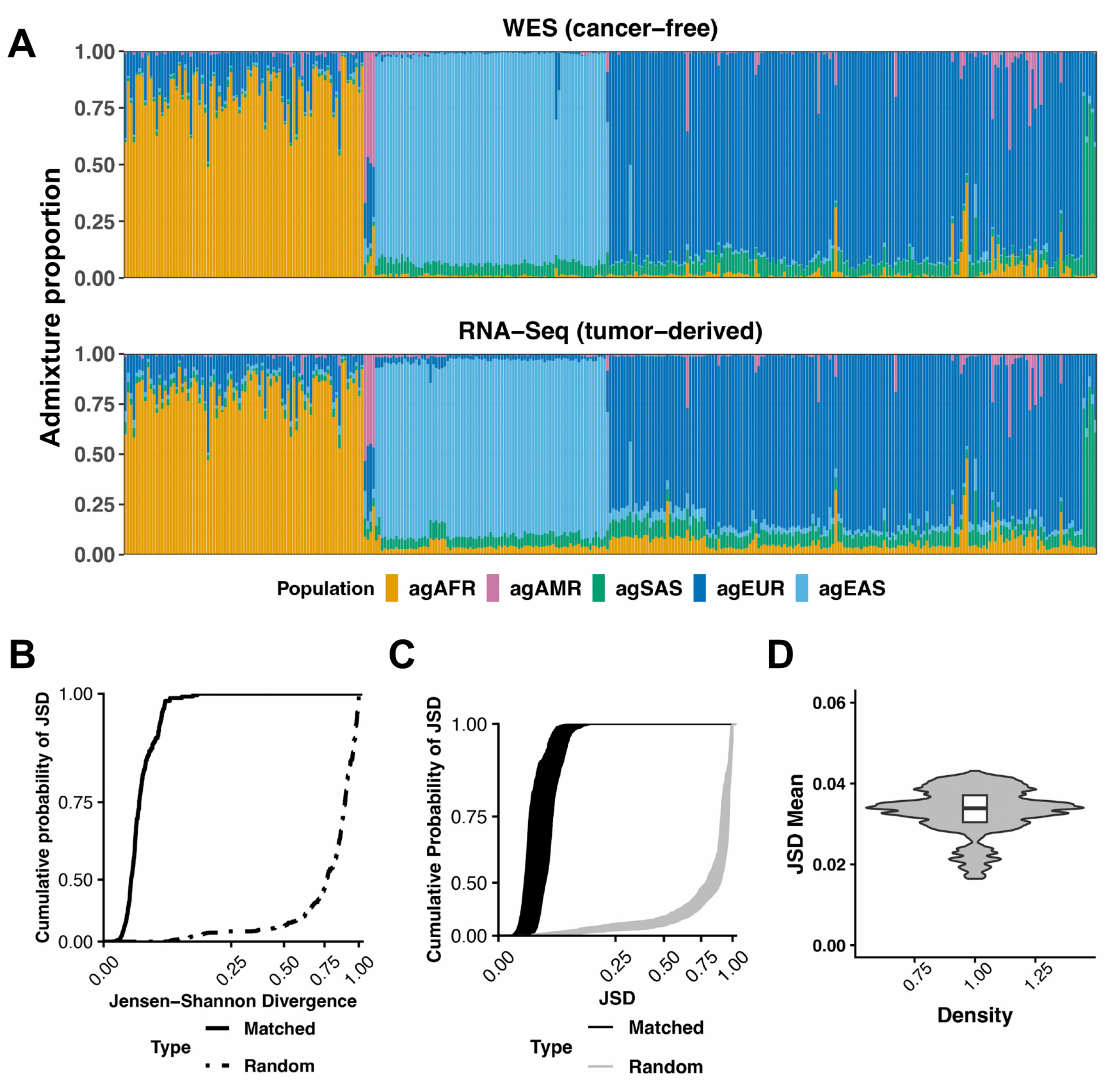


Figure S3: Analysis of admixture proportions, inferred from cancer-derived RNA-seq data in a cohort of 341 patient cases representing multiple cancer types, as indicated in Table S3, "Fig. S3 subset" column. A – Cohort-wide comparison of admixture proportions inferred from WES profiles of cancer-free DNA and matching RNA-Seq profiles of tumor-derived RNA. B – Cohort-wide cumulative distribution of pairwise JSD for admixture proportions inferred from cancer-derived RNA-Seq, with those from matching cancer-free whole-exome sequences. Cumulative JSD distribution for random pairs of patients is shown for reference. C – Cohort-wide family of cumulative distributions of the mean JSD measure of performance, estimated, for each patient, by data synthesis for each of the 341 cases of cancer-derived RNA. The corresponding family of distributions of JSD computed for random pairs of genotypes is shown for reference. D – Cohort-wide violin plot for the empirical mean JSD values, each estimated by data synthesis.

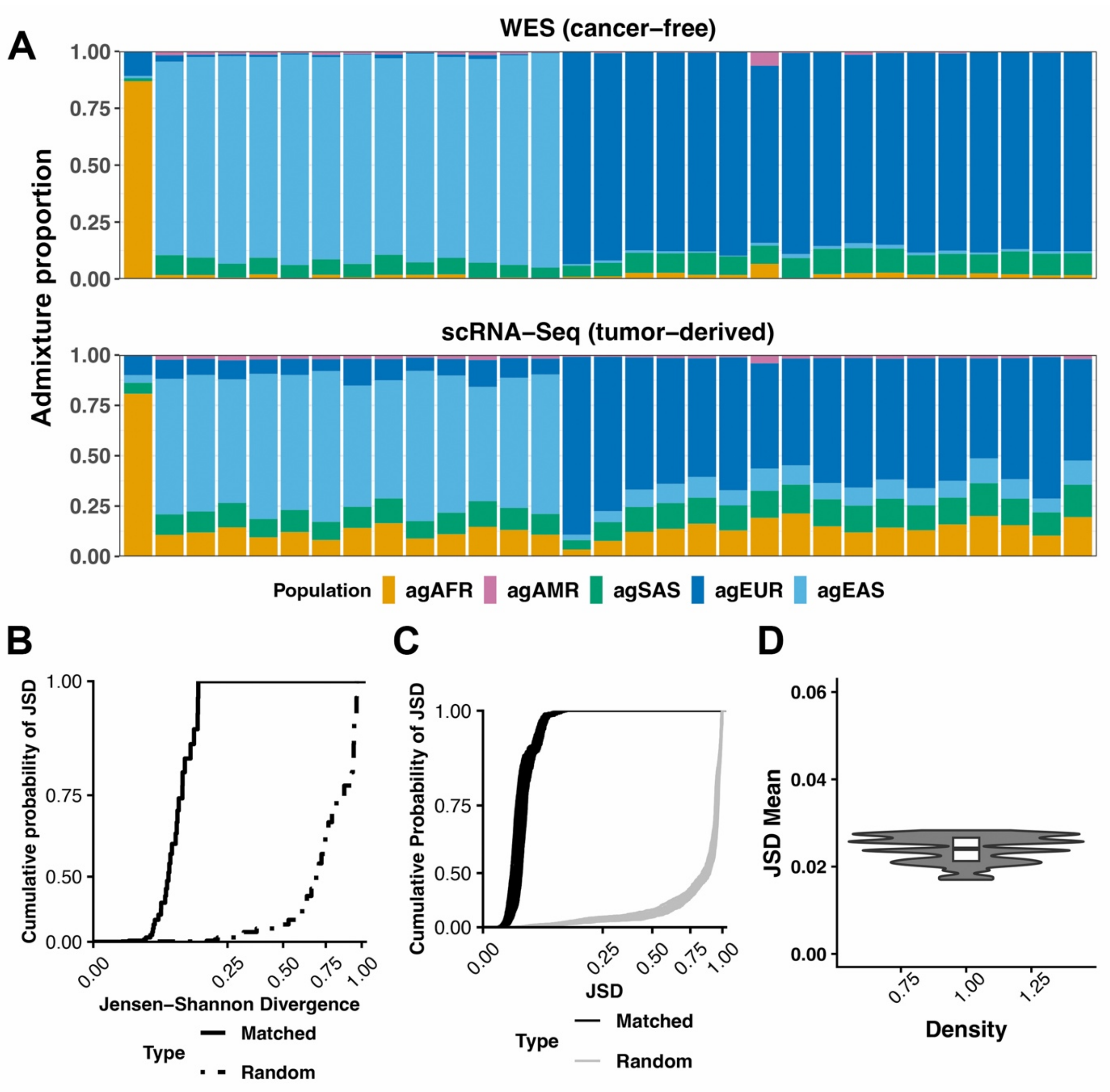


Figure S4: Analysis of admixture proportions, inferred from cancer-derived scRNA-Seq data in a cohort of 31 patient cases representing multiple cancer types, as indicated in Table S3, "Fig. S4 subset" column. A – Cohort-wide comparison of admixture proportions inferred from WES profiles of cancer-free DNA and matching scRNA-Seq profiles of tumor-derived scRNA. B – Cohort-wide cumulative distribution of pairwise JSD for admixture proportions inferred from cancer-derived scRNA-Seq, with those from matching cancer-free whole-exome sequences. Cumulative JSD distribution for random pairs of patients is shown for reference. C – Cohort-wide family of cumulative distributions of the mean JSD measure of performance, estimated, for each patient, by data synthesis for each of the 31 cases of cancer-derived scRNA. The corresponding family of distributions of JSD computed for random pairs of genotypes is shown for reference. D – Cohort-wide violin plot for the empirical mean JSD values, each estimated by data synthesis.

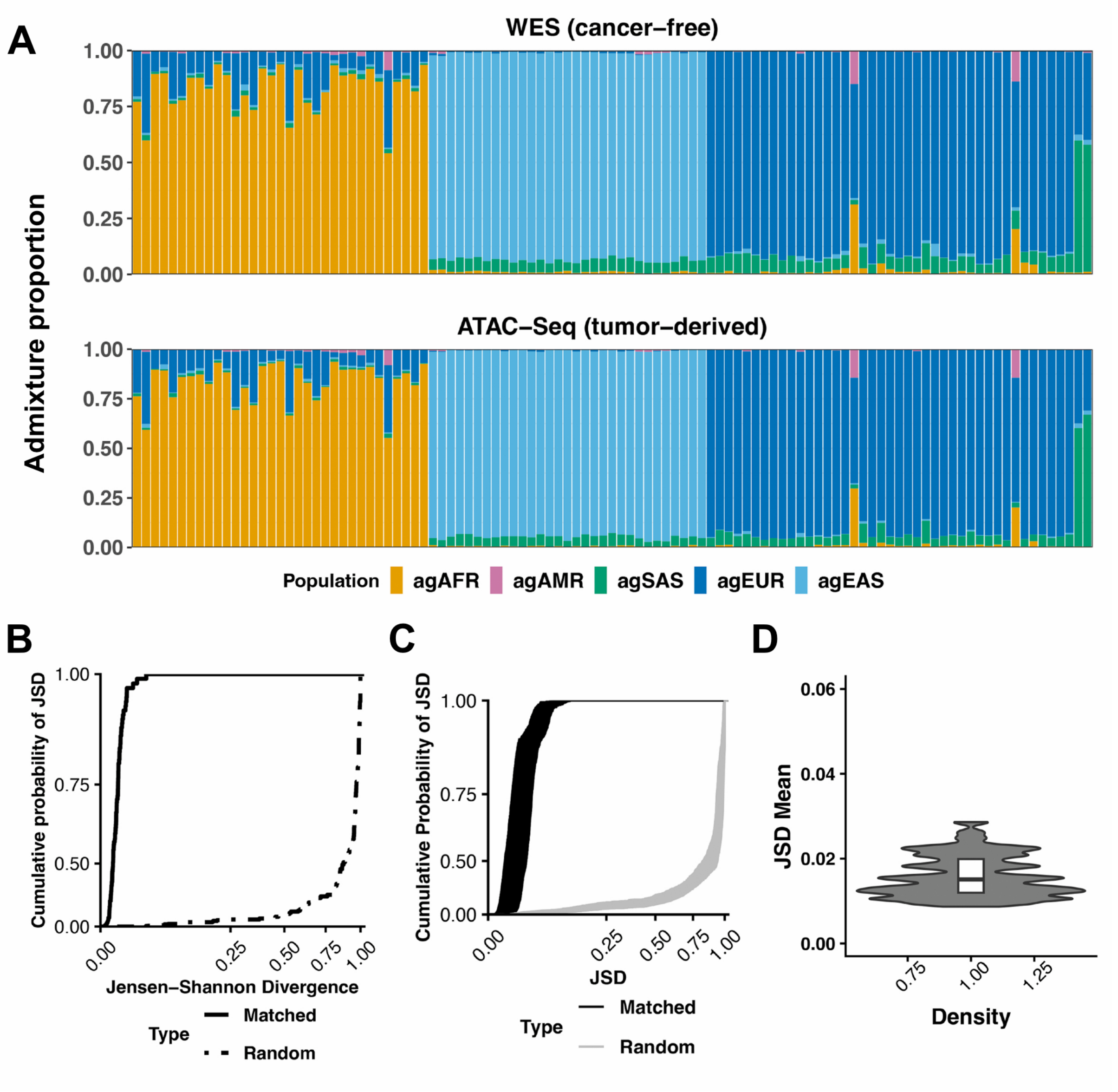


Figure S5: Analysis of admixture proportions, inferred from cancer-derived ATAC-Seq data in a cohort of 107 patient cases representing multiple cancer types, as indicated in Table S3, “Fig. S5 subset” column. A – Cohort-wide comparison of admixture proportions inferred from WES profiles of cancer-free DNA and matching ATAC-Seq profiles of tumor-derived DNA. B – Cohort-wide cumulative distribution of pairwise JSD for admixture proportions inferred from cancer-derived ATAC-Seq, with those from matching cancer-free whole-exome sequences. Cumulative JSD distribution for random pairs of patients is shown for reference. C – Cohort-wide family of cumulative distributions of the mean JSD measure of performance, estimated, for each patient, by data synthesis for each of the 107 cases of cancer-derived DNA. The corresponding family of distributions of JSD computed for random pairs of genotypes is shown for reference. D – Cohort-wide violin plot for the empirical mean JSD values, each estimated by data synthesis.

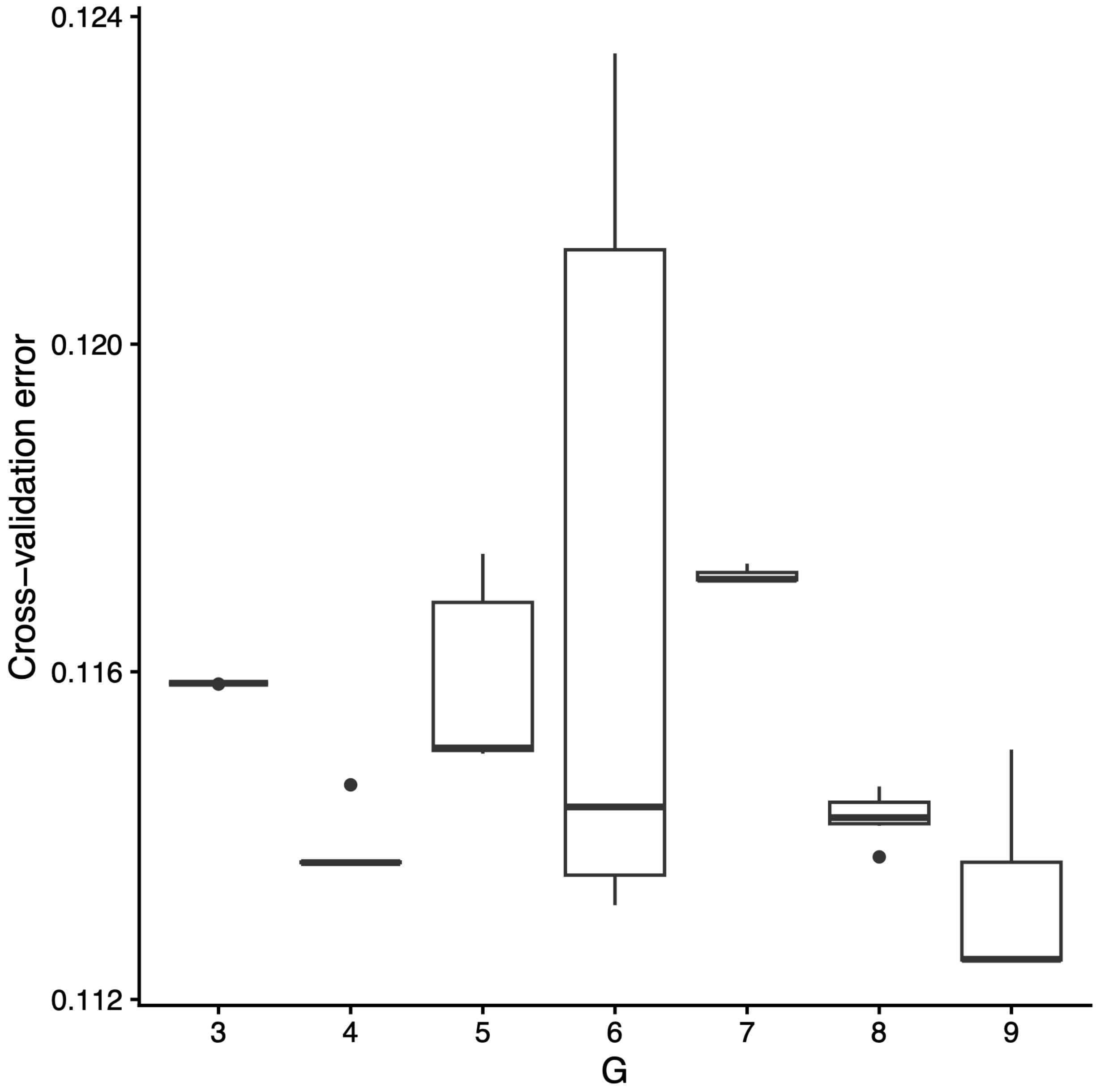


Figure S6: ADMIXTURE cross-validation error as a function of the number of ancestral groups G in the 1KG population-reference dataset.
