## Supplementary material for "A versatile set of computational tools for inference of genetic ancestry from human molecular data": tableCaptions.pdf

**Supplementary table captions**

S1: Summary of input molecular profile counts for datasets used in this work, with breakdown by clinical condition, tissue of origin, and experimental strategy.

S2: Detailed metadata for input data from repositories other than GDC, including clinical condition, tissue source, experimental strategy for molecular data of interest, experimental strategy for matching molecular data used for inference of ground truth, ground-truth ancestry, and self-identified race and ethnicity (SIRE).

S3: Donor metadata and ancestry findings for the GDC molecular data used in the study. The subsets of the GDC data as referred to in the Results section are as indicated in the respective columns.

S4: Tabulation of data underlying Figure 2A.

S5: Synthetic ChIP-Seq profiles and ancestry as a category inferred thereof (table.S5.csv.gz).

S6: Synthetic ATAC-Seq and scRNA-Seq profiles and ancestry as a category inferred thereof (table.S6.csv.gz).

S7: Tabulation of data underlying Figure 2B.

S8: Summary of the subset A — the 106 donors with tumor-derived data for all of WES, RNA-Seq and ATAC-Seq experimental strategies (WES, RNA-Seq, ATAC-Seq) and a matching cancer-free WES profile, underlying Figures 3 and 4 — organized by cancer type.

S9: Admixture proportions for all cancer-derived molecular profiles studied, those inferred from the matching cancer-free data, and the corresponding pairwise JSD values.

S10: Admixture proportions for all the profiles synthesized for inference of admixtures, the admixture proportions of the matching 1KG profile, and the JSD between them (table.S10.csv.gz).

S11: The ancestral composition of the 2,469 mutually unrelated individuals in the 1KG collection used as the population reference panel in this work.

S12: Software used to preprocess the input molecular profiles.
